# X-ray Structure of Main Protease of the Novel Coronavirus SARS-CoV-2 Enables Design of α-Ketoamide Inhibitors

**DOI:** 10.1101/2020.02.17.952879

**Authors:** Linlin Zhang, Daizong Lin, Xinyuanyuan Sun, Katharina Rox, Rolf Hilgenfeld

## Abstract

A novel coronavirus has been identified as the causative agent of a massive outbreak of atypical pneumonia originating at Wuhan, Hubei province, China. Involved in the formation of the coronavirus replication complex, the viral main protease (M^pro^, also called 3CL^pro^) represents an attractive target for therapy. We determined the crystal structure of the unliganded M^pro^ at 1.75 Å resolution and used this structure to guide optimization of a series of alpha-ketoamide inhibitors. The main goal of the optimization efforts was improvement of the pharmacokinetic properties of the compounds. We further describe 1.95- and 2.20-Å crystal structures of the complex between the enzyme and the most potent alpha-ketoamide optimized this way. These structures will form the basis for further development of these compounds to antiviral drugs.

## Introduction

Since December 2019, a new coronavirus has been emerging in the city of Wuhan, the capital of Hubei province in China. Whereas at the beginning of the outbreak, cases were connected to the Huanan seafood and animal market in Wuhan, efficient human-to-human transmission led to exponential growth in the number of cases, with the count standing at >71,000 as of today, with a ~2.4% case-fatality rate. The RNA genome of 2019-nCoV features an identity of about 82% to that of the SARS coronavirus (SARS-CoV); both viruses belong to clade b of the genus *Betacoronavirus*. Hence, it has been proposed to rename the new virus as SARS-CoV-2 (Gorbalenya et al., 2020).

One of the best characterized drug targets among coronaviruses is the main protease (M^pro^, also called 3CL^pro^). Along with the papain-like protease(s), this enzyme is essential for processing the polyproteins that are translated from the viral RNA (Hilgenfeld, 2014). The M^pro^ operates at no less than 11 cleavage sites on the large polyprotein 1ab (replicase 1ab, ~790 kDa); the recognition sequence at most sites is Leu - Gln↓(Ser,Ala,Gly) (↓ marks the cleavage site). Inhibiting the activity of this enzyme will block viral replication. Since no human proteases with a similar cleavage specificity are known, chances are that inhibitors will exhibit low toxicity.

Here we report the crystal structure of the M^pro^ of 2019-nCoV in the unliganded form and the usage of this structure to design new a-ketoamide inhibitors, compounds **13a** and **13b**. These compounds were developed on the basis of our first-generation peptidomimetic a-ketoamides, which we designed as broad-spectrum antivirals targeting the M^pro^s of beta-CoVs and alpha-CoVs, as well as the 3C protease of enteroviruses (Zhang et al., 2020). Compared to other inhibitors of cysteine proteases, a-ketoamides have the advantage of bearing a relatively mild electrophilic warhead that interacts with the catalytic center of the target enzyme through two hydrogen bonds rather than one, in addition to the covalent bond formed by nucleophilic attack of the catalytic cysteine onto the a-keto moiety.

## Results and Discussion

The 1.75-Å crystal structure of the unliganded 2019-nCoV M^pro^ (Fig. 1) reveals the high similarity to the SARS-CoV M^pro^ that was to be expected from sequence comparisons (Zhou et al., 2020; Wu et al., 2020). The r.m.s. deviation between the two free-enzyme structures is 0.53 Å for all Ca positions of the molecule but the isolated domains exhibit r.m.s. deviations of only 0.31 Å (domain II) - 0.42 Å (domains I, III) (comparison between the novel coronavirus M^pro^ structure and SARS-CoV M^pro^, PDB entry 2BX4 (Tan et al., 2005)). The chymotrypsin- and picornavirus 3C protease-like domains I and II (residues 10-99 and 100-182, respectively) are six-stranded antiparallel b-barrels that harbor the substrate-binding site between them. Domain III (residues 198-303), a globular cluster of five helices, is involved in the regulation of dimer formation of the M^pro^ (Shi & Song, 2006). 2019-nCoV M^pro^ forms a tight dimer (contact interface, predominantly between domain II of molecule A and the NH2-terminal residues of molecule B (“N-finger”): ~1394 Å^2^), with the two molecules oriented perpendicular to one another (Fig. 1). Dimerization of the enzyme is necessary for catalytic activity, because the N-finger of each of the two protomers interacts with Glu^166^ of the other protomer and thereby helps shape the S1 pocket of the substrate-binding site (Anand et al., 2002). To reach this interaction site, the N-finger is squeezed in between domains II and III of the parent monomer and domain II of the other monomer. Interestingly, in the SARS-CoV but not in the 2019-nCoV M^pro^ dimer, there is a polar interaction between the two domains III involving a 2.60-Å hydrogen bond between the side-chain hydroxyl groups of residue Thr^285^ of each protomer, and supported by a hydrophobic contact between the side-chain of Ile^286^ and Thr^285^ Cg2. In 2019-nCoV, the threonine is replaced by alanine (indicated by the black sphere in Fig. 1), and the isoleucine by leucine. It has previously been shown that replacing Ser^284^, Thr^285^, and Ile^286^ by alanine residues in SARS-CoV M^pro^ leads to a 3.6-fold enhancement of the catalytic activity of the protease, concomitant with a slightly closer packing of the two domains III of the dimer against one another (Lim et al., 2014). According to that study, this is accompanied by changes of the structural dynamics of the enzyme that transmit the effect of the mutation to the catalytic center. Indeed, the Thr^285^Ala replacement observed in the 2019-nCoV M^pro^ also allows the two domains III to approach each other a little closer (the distance between the Ca atoms of residues 285 in molecules A and B is 6.77 Å in SARS-CoV M^pro^ and 5.21 Å in 2019-nCoV M^pro^ and the distance between the centers of mass of the two domains III shrinks from 33.4 Å to 32.1 Å).

**Fig. 1:**
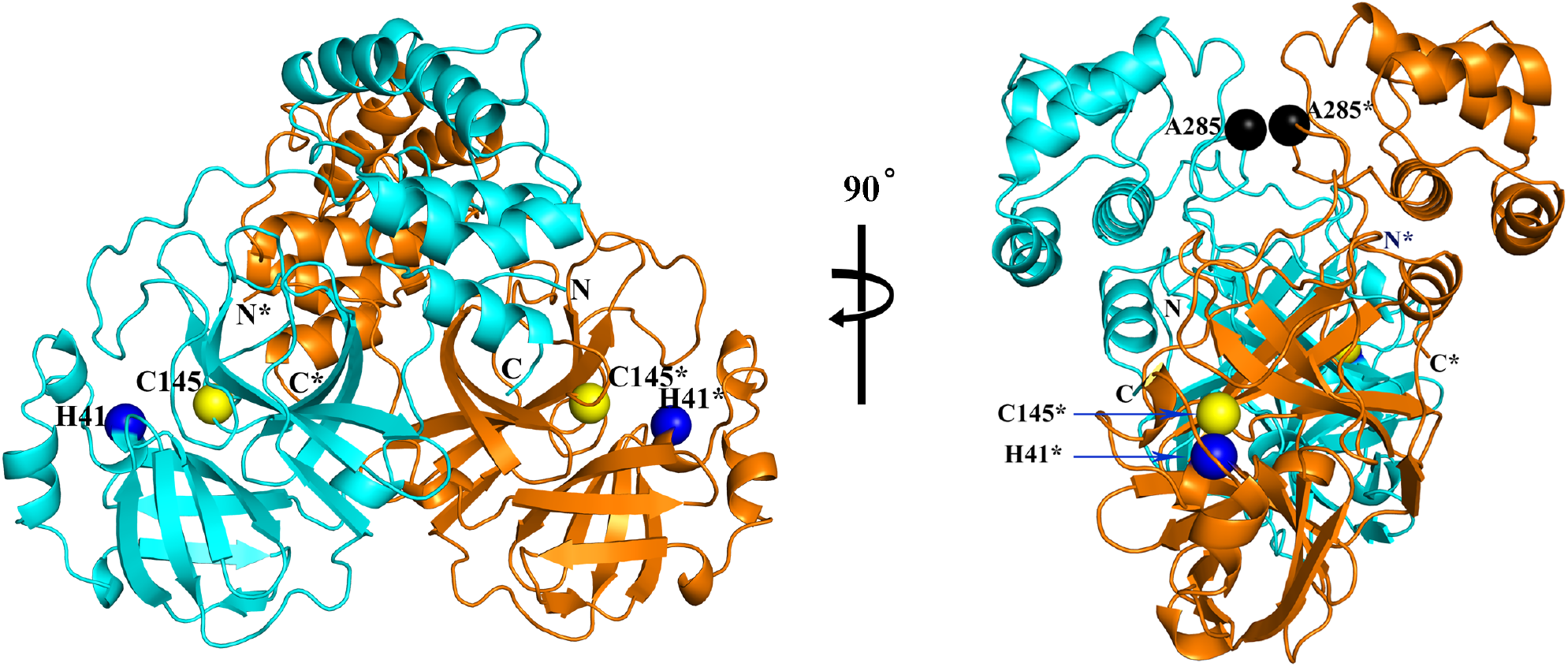
**Three-dimensional structure of 2019-nCoV M^pro^**, in two different views. One protomer of the dimer is shown in light blue, the other one in orange. Amino-acid residues of the catalytic site are indicated as yellow and blue spheres, for Cys^145^ and His^41^, respectively. Black spheres indicate the positions of Ala^285^ of each of the two domains III (see text). Chain termini are labeled N and C for molecule A (light blue) and N* and C* for molecule B (orange).

In the active site of 2019-nCoV M^pro^, Cys^145^ and His^41^ form a catalytic dyad. Like in SARS-CoV M^pro^ and other coronavirus homologues, a buried water molecule is found hydrogen-bonded to His^41^; this could be considered the third component of a catalytic triad.

Previously, we have designed and synthesized peptidomimetic a-ketoamides as broadspectrum inhibitors of the main proteases of betacoronaviruses and alphacoronaviruses as well as the 3C proteases of enteroviruses (Zhang et al., 2020). The best of these compounds (**11r**; see Scheme 1) showed an EC50 of 400 picomolar against MERS-CoV in Huh7 cells as well as low micromolar EC50 values against SARS-CoV and a whole range of enteroviruses in various cell lines. In order to improve the half-life and the solubility of the compounds in human plasma, and to reduce the binding to plasma proteins, we have modified the compound by hiding the P3 - P2 amide bond within a pyridone ring and by replacing the cinnamoyl group. For a compound related to **11r** but modified this way (compound **13a**), the half-life in human plasma was increased by 50%, solubility was improved, and plasma protein binding was reduced from 99% to 94%. There was no sign of toxicity in mice. In addition, **13a** showed good metabolic stability using mouse and human microsomes, with intrinsic clearance rates Cl_int_mouse_= 32.00 μL/min/mg protein and Cl_int_human_= 20.97 μL/min/mg protein. This means that after 30 min, around 80% for mouse and 60% for humans, respectively, of residual compound remained metabolically stable. Pharmacokinetic studies in CD-1 mice using the subcutaneous route at 20 mg/kg showed that **13a** stayed in plasma for up to only 4 hrs, but was excreted via urine up to 24 hrs. The Cmax was determined at 334.50 ng/mL and the mean residence time was about 1.59 hrs. Although **13a** seemed to be cleared very rapidly from plasma, it was found at 24 hrs at 135 ng/g tissue in the lung and at 52.7 ng/mL in broncheo-alveolar lavage fluid (BALF) suggesting that it was mainly distributed to tissue. In the light of the current CoV outbreak, it is advisable to develop compounds with lung tropism such as **13a.** However, compared to **11r,** the structural modification led to some loss of inhibitory activity against the main protease of 2019-nCoV (IC50 = 2.39 ± 0.63 uM) as well as the 3C proteases of enteroviruses. To enhance the antiviral activity against betacoronaviruses of clade b (2019-nCoV and SARS-CoV), we sacrificed the goal of broad-spectrum activity including the enteroviruses for the time being and replaced the P2 cyclohexyl moiety of **13a** by cyclopropyl in **13b**, because the S2 pocket of the betacoronavirus main proteases shows a pronounced plasticity enabling it to adapt to the shape of smaller inhibitor moieties entering this site (Zhang et al., 2020). Here we present X-ray crystal structures in two different crystal forms, at 1.95 and 2.20 Å resolution, of the complex between α-ketoamide **13b** optimized this way and the M^pro^ of 2019-nCoV (Fig. 2). One structure is in space group *C*2, where both protomers of the M^pro^ dimer are bound by crystal symmetry to have identical conformations, the other is in space group *P*2_1_2_1_2_1_, where the two protomers are independent of each other and free to adopt different conformations. Indeed, we find that in the latter crystal structure, the key residue Glu^166^ adopts an inactive conformation (as evidenced by its prolonged distance from His^172^ and the lack of H-bonding interaction between Glu^166^ and the P1 moiety of the inhibitor (see below)), even though compound **13b** is bound in the same mode as in molecule A. This phenomenon has also been observed, in a more pronounced form, with the SARS-CoV M^pro^ (Yang et al., 2003) and is consistent with the half-site activity described for this enzyme (Chen et al., 2006). In all copies of the inhibited 2019-nCoV M^pro^, the inhibitor binds to the shallow substrate-binding site at the surface of each protomer, between domains I and II (Fig. 2).

**Scheme 1:**
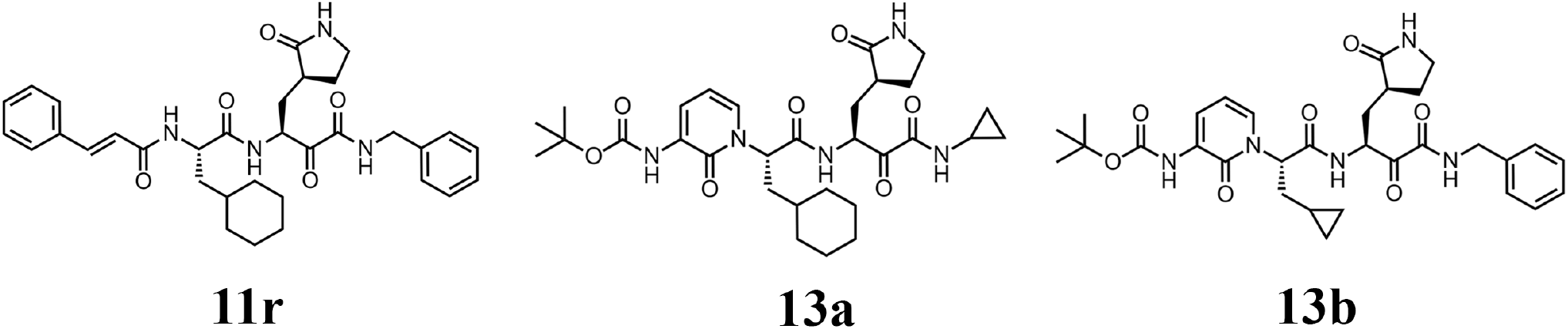
Chemical structures of α-ketoamide inhibitors **11r**, **13a**, and **13b**

**Fig. 2:**
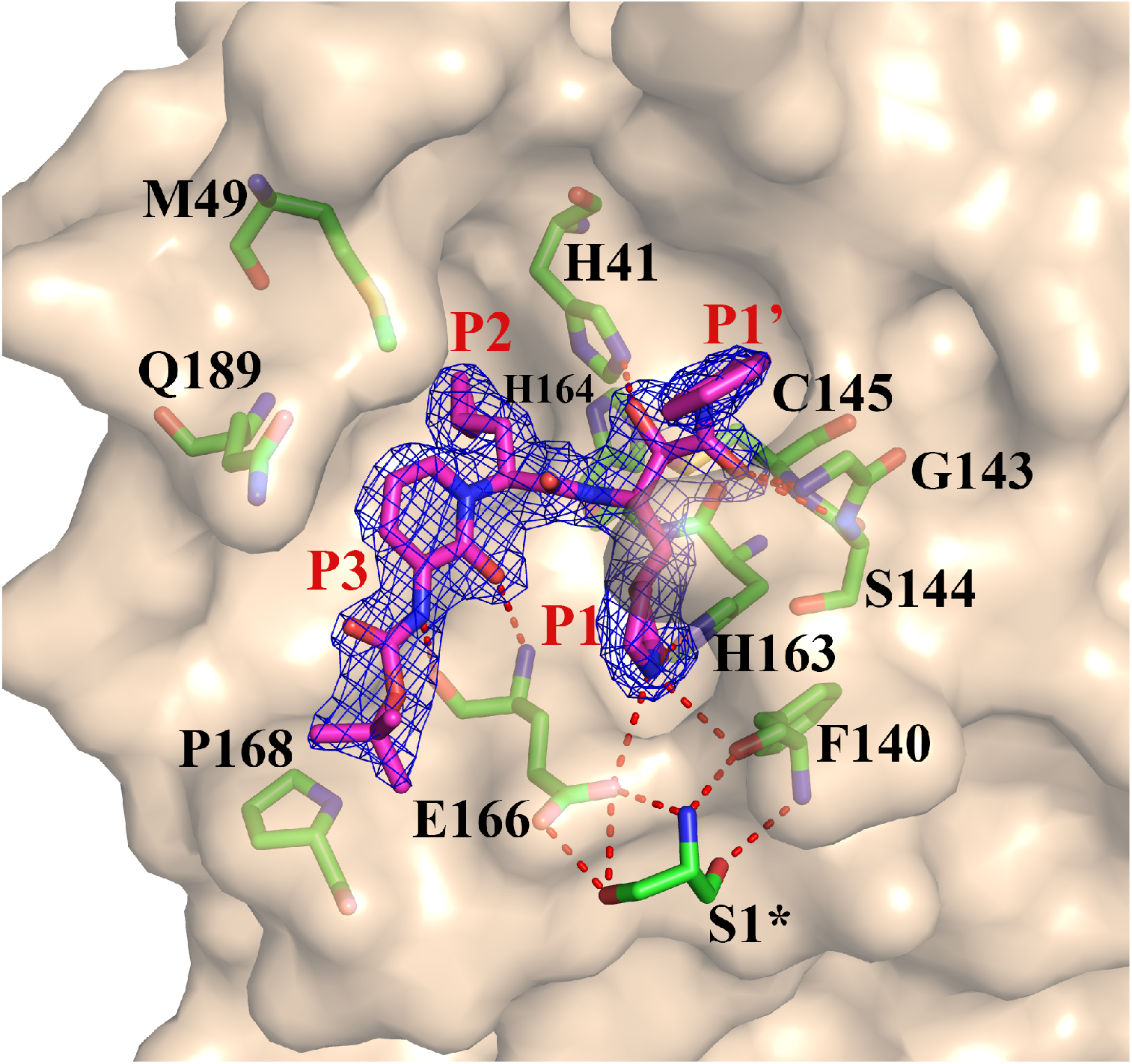
**Compound 13b in the substrate-binding cleft located between domains I and II of the M^pro^**, in the monoclinic crystal form (space group *C*2). 2F_o_-F_c_ electron density is shown for the inhibitor (contouring level: 1σ). Carbon atoms of the inhibitor are magenta, oxygens red, nitrogens blue, and sulfur yellow. Note the interaction between the N-terminal residue of chain B, S^1^*, and E^166^ of chain A.

Through the nucleophilic attack of the catalytic Cys^145^ onto the a-keto group of the inhibitor, a thiohemiketal is formed in a reversible reaction. This is clearly reflected in the electron density (Fig. 2); the stereochemistry of this chiral moiety is *S* in all three copies of compound **13b** in these structures. The oxyanion (or hydroxyl) group of this thiohemiketal is stabilized by a hydrogen bond from His^41^, whereas the amide oxygen of **13b** accepts a hydrogen bond from the main-chain amides of Gly^143^, Cys^145^, and partly Ser^144^, which form the canonical “oxyanion hole” of the cysteine protease. The P1 g-lactam moiety, designed as a glutamine surrogate (Dragovich et al., 1999; Tan et al., 2013), is deeply embedded in the S1 pocket of the protease, where the lactam amide nitrogen donates a three-center (bifurcated) hydrogen bond to the main-chain oxygen of Phe^140^ (3.20/3.10/3.28 Å; values for the structure in space group *C*2/space group *P*2_1_2_1_2_1_ molecule A/space group *P*2_1_2_1_2_1_ molecule B) and to the Glu^166^ carboxylate (3.35/3.33/ - Å), and the carbonyl oxygen accepts a 2.57/2.51/2.81-Å H-bond from the imidazole of His^163^. The P2 cyclopropyl methyl moiety is embraced by the S2 subsite, which has shrunk by 28.4 Å^3^ compared to a complex between compound **13a** with P2 = cyclohexyl methyl and the SARS-CoV M^pro^ (LLZ et al., unpublished). The pyridone in the P3 position of the inhibitor occupies the space normally filled by the substrate’s main chain, its carbonyl oxygen accepts a 2.89/2.99/3.00-Å hydrogen bond from the main-chain amide of residue Glu166. Further, the P3 amide donates a 2.83/2.96/2.87-Å H-bond to the main-chain oxygen of Glu^166^. Embedded within the pyridone, the P2 nitrogen can no longer donate a hydrogen bond to the protein; however, our previous crystal structures showed that the P2 main-chain amide of the linear α-ketoamides does not make a hydrogen bond with the protein in all cases, so this interaction does not seem to be essential (Zhang et al., 2020). The Boc group does not occupy the canonical S4 site of the protease, but is located near Pro^168^ (at a distance of 3.81/4.17/3.65 Å); due to this interaction, the latter residue is “pushed” by >2 Å (compared to the structure of the free enzyme). This contact explains that removing the Boc group weakens the inhibitory potency of this compound by a factor of about 2. Interestingly, there is a space between the pyridone ring of **13b**, the main chain of residue Thr^190^, and the side-chain of Gln^189^, which is filled by a DMSO molecule in the *C*2 crystal structure and a water molecule in the *P*2_1_2_1_2_1_ structure. This suggests that P3 moieties more bulky than pyridone may be accepted here.

Compound **13b** inhibits the purified recombinant 2019-nCoV M^pro^ with IC_50_ = 0.67 ± 0.18 μM. The corresponding IC50 values for inhibition of the SARS-CoV M^pro^ and the MERS-CoV M^pro^ are 0.9 ± 0.29 μM and 0.58 ± 0.22 μM, respectively. In a SARS-CoV replicon (Kusov et al., 2015), RNA replication is inhibited with EC50 = 1.75 ± 0.25 μM. In human Calu3 cells infected with MERS-CoV, the compound showed excellent antiviral activity (Lucie Sauerhering & Stephan Becker personal communication).

As the next steps in the development of compound **13b** towards a potential drug targeting 2019-nCoV or other coronaviruses, we will undertake tests in virus-infected cell cultures and in a small-animal model (once it will become available for 2019-nCoV). Meanwhile, our crystal structures may be used by others for virtual screening and *de-novo* design of inhibitors.

## Acknowledgements

The authors are grateful to Yuri Kusov and Guido Hansen as well as Aws Aljnabi for determining the inhibitory activities of compounds in a SARS-CoV replicon and against recombinant MERS-CoV M^pro^, respectively. We are indebted to Andrea Ahlers and Janine Schreiber for excellent technical assistance and to the staff at beamLine 14.2 of BESSY II, Berlin, Germany, for their support during diffraction data collection. We thank the German Center for Infection Research (DZIF) for financial support (projects TTU01.806 and TTU09.710). Crystallographic coordinates and structure factors are available from the PDB under accession codes 6Y2E (unliganded M^pro^), 6Y2F (complex with **13b** in space group *C*2), and 6Y2G (complex with **13b** in space group *P*2_1_2_1_2_1_). The plasmid encoding the 2019-nCoV M^pro^ will be freely available. The available amounts of compounds **13a** and **13b** are limited.

## References

Anand K., Palm G. J., Mesters J. R., Siddell S. G., Ziebuhr J., Hilgenfeld R., Structure of coronavirus main proteinase reveals combination of a chymotrypsin fold with an extra alpha-helical domain. EMBO J. 21, 3213–3224 (2002).

Chen H., Wei P., Huang C., Tan L., Liu Y., Lai L., Only one protomer is active in the dimer of SARS 3C-like proteinase. J. Biol. Chem. 281, 13894–13898 (2006).

Dragovich P. S., Zhou R., Skalitzky D. J., Fuhrman S. A., Patick A. K., Ford C. E., Meador, 3rd J. W., Worland S. T., Solid-phase synthesis of irreversible human rhinovirus 3C protease inhibitors. Part 1: Optimization of tripeptides incorporating N-terminal amides. Bioorg. Med. Chem. 7, 589–598 (1999).

Gorbalenya A. E., Baker S. C., Baric R. S., de Groot R. J., Drosten C., Gulyaeva A. A., Haagmans B. L., Lauber C, Leontovich A. M., Neuman B. W., Penzar D., Perlman S., Poon L. L. M., Samborskiy D., Sidorov I. A., Sola I., Ziebuhr J., Severe acute respiratory syndrome-related coronavirus: The species and its viruses – a statement of the Coronavirus Study Group. biorxiv preprint server (2020) doi: https://doi.org/10.1101/2020.02.07.937862.

Hilgenfeld R., From SARS to MERS: Crystallographic studies on coronaviral proteases enable antiviral drug design. FEBS J. 281, 4085–4096 (2014).

Kusov Y., Tan J., Alvarez E., Enjuanes L., Hilgenfeld R., A G-quadruplex-binding macrodomain within the “SARS-unique domain” is essential for the activity of the SARS-coronavirus replication-transcription complex. Virology 484, 313–322 (2015).

Lim L., Shi J., Mu Y., Song J., Dynamically-driven enhancement of the catalytic machinery of the SARS 3C-like protease by the S284-T285-I286/A mutations on the extra domain. PLoS One 9, e101941 (2014).

Shi J., Song J., The catalysis of the SARS 3C-like protease is under extensive regulation by its extra domain. FEBS J. 273, 1035–1045 (2006).

Tan J., George S., Kusov Y., Perbandt M., Anemuller S., Mesters J. R., Norder H., Coutard B., Lacroix C., Leyssen P., Neyts J., Hilgenfeld R., 3C protease of enterovirus 68: structure-based design of Michael acceptor inhibitors and their broad-spectrum antiviral effects against picornaviruses. J. Virol. 87, 4339–4351 (2013).

Tan J., Verschueren K. H. G., Anand K., Shen J., Yang M., Xu Y., Rao Z., Bigalke J., Heisen B., Mesters J. R., Chen K., Shen X., Jiang H., Hilgenfeld R., pH-dependent conformational flexibility of the SARS-CoV main proteinase (M^pro^) dimer: Molecular dynamics simulations and multiple X-ray structure analyses. J. Mol. Biol. 354, 25–40 (2005).

Wu F., Zhao S., Yu B., Chen Y.-M., Wang W., Song Z.-G., Hu Y., Tao Z.-W., Tian J.-H., Pei Y.-Y., Yuan M.-L., Zhang Y.-L., Dai F.-H., Liu Y., Wang Q.-M., Zheng J.-J., Xu L., Holmes E. C., Zhang Y.-Z., A new coronavirus associated with human respiratory disease in China. Nature https://doi.org/10.1038/s41586-020-2008-3 (2020).

Yang H., Yang M., Ding Y., Liu Y., Lou Z., Zhou Z., Sun L., Mo L., Ye S., Pang H., Gao G. F., Anand K., Bartlam M., Hilgenfeld R., Rao Z., The crystal structures of severe acute respiratory syndrome virus main protease and its complex with an inhibitor. Prac. Natl. Acad. Sci. USA 100, 13190–13195 (2003).

Zhang L., Lin D., Kusov Y., Nian Y., Ma Q., Wang J., von Brunn A., Leyssen P., Lanko K., Neyts J., de Wilde A., Snijder E. J., Liu H., Hilgenfeld R, Alpha-ketoamides as broad-spectrum inhibitors of coronavirus and enterovirus replication: Structure-based design, synthesis, and activity assessment. J. Med. Chem., in press (2020). Biorxiv preprint server, doi: 10.1101/2020.02.10.936898.

Zhou P., Yang X.-L., Wang X.-G., Hu B., Zhang L., Zhang W., Si H.-R., Zhu Y., Li B., Huang C.-L., Chen H.-D., Chen J., Luo Y., Guo H., Jiang R.-D., Liu M.-Q., Chen Y., Shen X.-R., Wang X., Zheng X.-S., Zhao K., Chen Q.-J., Deng F., Liu L.-L., Yan B., Zhan F.-X., Wang Y.-Y., Xiao G.-F., Shi Z.-L., A pneumonia outbreak associated with a new coronavirus of probable bat origin. Nature https://doi.org/10.1038/s41586-020-2012-7 (2020).

