## Supplementary materials for "X-ray Structure of Main Protease of the Novel Coronavirus SARS-CoV-2 Enables Design of α-Ketoamide Inhibitors"

### Materials & Methods

#### Recombinant protein production

A gene encoding 2019-nCoV $M^{pro}$ (ORF1ab polyprotein residues 3264-3569, GenBank code: MN908947.3) with *Escherichia coli* codon usage was synthesized by MWG Eurofins. The synthesized gene was first amplified with the forward primer 5′-TCGGGGTTTCGCAAAAT-3′ and the reverse primer 5′-CTGAAACGTGACACCGCTACA -3′. Subsequently, this PCR product was employed as the template for the amplification of the target gene with primers (forward) 5′-CGCGGATCCTCGGCAGTGCTGCAATCGGGGTTTCGCAAAAT-3′, and (reverse) 5′-CCGCTCGAGTTAATGATGATGATGATGATGGGGTCCCTGAAACGTGACACCGCT ACACT -3′ including the cleavage sites of restriction enzymes for cloning into the vector PGEX-6p-1 (GE Healthcare). The amplified PCR product was digested with *BamHI* and *XhoI* and ligated into the vector PGEX-6p-1 digested with the same restriction enzymes. At the N-terminus, the construct designed for 2019-nCoV $M^{pro}$ contains the $M^{pro}$ cleavage-site (SAVLQ↓SGFRK; arrow indicates the cleavage site) corresponding to the cleavage site between Nsp4 and Nsp5 in the polyprotein of this virus. At the C-terminus, the construct codes for a modified PreScission cleavage site (SGVTFQ↓GP; the six-residue sequence at the C-terminus of the 2019-nCoV $M^{pro}$ was used as P6-P1 for PreScission cleavage) connected to a $His_6$-tag. An authentic N-terminus was generated during gene expression by auto-cleavage of the $M^{pro}$ itself, and the authentic C-terminus was generated after the treatment with PreScission protease, similar to the approach described for SARS-CoV $M^{pro}$ in (Xue et al., 2007). The gene sequence of the $M^{pro}$ was verified by sequencing at MWG Eurofins.

The sequence-verified 2019-nCoV $M^{pro}$ construct was transformed into *E. coli* strain BL21-Gold (DE3) (Novagen). Transformed clones were pre-cultured at 37°C in 50 mL 1 x YT medium with ampicillin (100 μg/mL) for 3 hrs, and the incubated culture was inoculated into 4 L 1 x YT medium supplied with 100 μg/mL ampicillin. 0.5 mM isopropyl-D-thiogalactoside (IPTG) was added for induction of the overexpression of the $M^{pro}$ gene at 37°C when the $OD_{600}$ reached 0.8. After 5 hrs, cells were harvested by centrifugation at 9954 x *g*, 4°C for 15 min. The pellets were resuspended in 30 mL buffer A (20 mM Tris, 150 mM NaCl, pH 7.8; pH of all buffers was adjusted at room temperature) and then lysed by sonication on ice. The lysate was clarified by ultracentrifugation at 146,682 x *g* at 4°C for 1 h. The supernatant was loaded onto a HisTrap FF column (GE Healthcare) equilibrated with buffer A. The HisTrap FF column was washed with 150 mL buffer A to remove unspecific binding proteins, followed by elution using buffer B (150 mM NaCl, 20 mM Tris, 500 mM imidazole, pH 7.8) with a linear gradient of imidazole ranging from 0 mM to 500 mM, 20 column volumes. The fractions containing target protein were pooled and mixed with PreScission protease at a molar ratio of 5:1 and dialyzed into buffer C (20 mM Tris, 150 mM NaCl, 1 mM DTT, pH 7.8) at 4°C overnight, resulting in the target protein with authentic N- and C-termini. The PreScission-treated $M^{pro}$ was applied to the connected GSTtrap FF (GE Healthcare) and nickel columns to remove the GST-tagged PreScission protease, the His-tag, and protein with uncleaved His-tag. The His-tag-free $M^{pro}$ in the flow-through was loaded onto a HiTrap Q FF column (GE Healthcare) equilibrated with buffer D (20 mM Tris, 1 mM DTT, pH 8.0) for further purification. The column was eluted by buffer E (20 mM Tris, 1 M NaCl, 1 mM DTT, pH 8.0) with a linear gradient ranging from 0 to 500 mM NaCl (20 column volumes buffer). Fractions eluted from the Hitrap Q FF column containing the target protein with high purity were pooled and subjected to buffer exchange (20 mM Tris-HCl, 150 mM NaCl, 1 mM EDTA, 1 mM DTT, pH 7.8) by using Amicon Ultra 15 centrifugal filters (10 kD, Merck Millipore) at 2773 x *g*, and 4°C.

**Crystallization of the free 2019-nCoV $M^{pro}$**

A freshly prepared protein solution at a concentration of 25 mg/mL was cleared by centrifugation at 12,000 x *g*. Subsequently, a basic screen with the commercially available screening kit PACT premier™ HT-96 (Molecular Dimensions) was performed by using a Gryphon LCP crystallization robot (Art Robbins) employing the sitting-drop vapor-diffusion method at 18°C. 0.15 µL of protein solution and 0.15 µL of reservoir were mixed to equilibrate against 40 µL reservoir solution. Crystals appeared overnight under several conditions, e.g. condition B5 (0.1 M MIB (sodium malonate, imidazole, and boric acid in molar ratio 2:3:3), pH 8.0, 25% polyethylene glycol (PEG) 1,500), condition D4 (0.1 M MMT (DL-malic acid, MES and Tris base in molar ratio 1:2:2), pH 7.0, 25% PEG 1,500), condition E9 (0.2 M potassium sodium tartrate tetrahydrate, 20% PEG 3,350), etc. Crystals were fished from the drops and cryo-protected by mother liquor plus varied concentrations of glycerol (10%-20%). Subsequently, fished crystals were flash-cooled in liquid nitrogen.

**Crystallization of 13b in complex with 2019-nCoV $M^{pro}$**

Freshly prepared protein (as described above) at a concentration of 25 mg/mL was mixed with **13b** (dissolved in 100% DMSO) at a molar ratio of 1:5. The mixture was incubated at

4°C overnight. The next day, centrifugation was applied (12,000 x *g*) to remove the white precipitant. Subsequently, the supernatant was subjected to crystallization screening using the same method as described above for the crystallization of the free enzyme. A basic screen was applied by using three commercially available kits: PEGRx$^{TM}$ 1 & 2 (Hampton Research), PACT premier$^{TM}$ HT-96 (Molecular Dimensions), and Morpheus HT-96 (Molecular Dimensions). Crystals appeared overnight under conditions No. 45 (0.1 M bicine, pH 8.5, 20% PEG 10,000) of PEGRx$^{TM}$ 1 and No. 39 (10% PEG 200, 0.1 M bis-tris propane, pH 9.0, 18% PEG 8,000) of PEGRx$^{TM}$ 2, condition G9 (0.1 M Carboxylic acids (0.2 M sodium formate, 0.2 M ammonium acetate, 0.2 M sodium citrate tribasic dihydrate, 0.2 M potassium sodium tartrate tetrahydrate, 0.2 M sodium oxamate), 0.1 M buffer system 3 (1.0 M Tris (base), bicine, pH 8.5), pH 8.5, 30% precipitant mix 1 (20% v/v PEG 500 methyl ether, 10% PEG 20,000)) of Morpheus HT-96. Manual reproduction was performed by mixing 0.5 μL of complex and 0.5 μL of reservoir, equilibrating against 40 μL of reservoir in a 96-well plate under condition No. 45 of PEGRx$^{TM}$ 1 and No. 39 of PEGRx$^{TM}$ 2, No. G9 of Morpheus HT-96, by using the sitting-drop vapor-diffusion method. Crystals appeared also overnight in the manually reproduced drops. Crystals were fished from the reproduced condition-45 drop (cryo-protectant: mother liquor plus 15% glycerol, and 5 mM **13b**), the reproduced condition-39 drop (cryo-protectant: mother liquor plus 5 mM **13b**), and the original condition-G9 drop (cryo-protectant: mother liquor plus 5 mM **13b**). Subsequently, fished crystals were flash-cooled in liquid nitrogen.

**Diffraction data collection, phase determination, model building, and refinement**

All diffraction data sets were collected using synchrotron radiation of wavelength 0.9184 Å at beamLine BL14.2 of BESSY (Berlin, Germany), using a Pilatus3S 2M detector (Dectris) (Mueller et al., 2012). The data set of the free 2019-nCoV $M^{pro}$ was collected from a crystal grown under condition No. D4 of the PACT premier™ HT-96 in the basic screen 96-well plate. Two data sets of the 2019-nCoV $M^{pro}$ in complex with **13b** were collected from the crystals grown under condition No. 39 of PEGRx™ 2 (manual optimization) and No. G9 of Morpheus HT-96, respectively. *XDSapp* (Krug & Weiss, 2012), *Pointless* (Evans, 2006; Evans, 2011), and *Scala* (Evans, 2006) (the latter two from the CCP4 suite (Winn et al., 2011)) were used for processing the datasets. The diffraction dataset of the free 2019-nCoV $M^{pro}$ was processed at a resolution of 1.75 Å, in space group *C*2 (Table S2). For the complex of 2019-nCoV $M^{pro}$ with **13b**, the dataset collected from a condition-39 crystal was processed at a resolution of 1.95 Å and in space group *C*2 (Table S2), whereas the dataset collected from a condition-G9 crystal extended to a Bragg spacing of 2.20 Å, in space group $P2_12_12_1$ (Table S2). All three structures were determined by molecular replacement with the crystal structure of the free enzyme of the SARS-CoV $M^{pro}$ (PDB entry 2BX4 (Tan et al., 2005)) as search model, using the *Molrep* program (Winn et al., 2011; Vagin & Teplyakov, 2010). *Jligand* from the CCP4 suite (Winn et al., 2011; Lebedev et al., 2012) was employed for the generation of the geometric restraints for **13b**, and the inhibitor was built into the $F_o$-$F_c$ density by using the *Coot* software (Emsley et al., 2010). Refinement of the three structures was performed with *Refmac5* (Winn et al., 2011; Murshudov et al., 2011). Statistics of diffraction data processing and the model refinement are given in Table S2.

**Determination of inhibition of $M^{pro}$ by 13a and 13b**

A fluorescent substrate harboring the cleavage site (indicated by the arrow) of 2019-nCoV $M^{pro}$ (Dabcyl-KTSAVLQ↓SGFRKM-E(Edans)-$NH_2$), and buffer composed of 20 mM Tris, 100 mM NaCl, 1 mM EDTA, 1 mM DTT, pH 7.3 was used for the inhibition assay. In the fluorescence resonance energy transfer (FRET)-based cleavage assay (Zhu et al., 2011), the fluorescence signal of the Edans generated due to the cleavage of the substrate by the $M^{pro}$ was monitored at an emission wavelength of 460 nm with excitation at 360 nm, using a Flx800 fluorescence spectrophotometer (BioTek). Stock solutions of the compounds were prepared with 100% DMSO. For the determination of the $IC_{50}$, 0.5 μM of 2019-nCoV $M^{pro}$ was incubated with **13a** or **13b** at various concentrations from 0 to 100 μM in reaction buffer at 37°C for 10 min. Afterwards, the FRET substrate at a final concentration of 20 μM was added to each well at a final total volume of 50 μL to initiate the reaction. The GraphPad Prism 6.0 software (GraphPad) was used for the calculation of the $IC_{50}$ values. Measurements of inhibitory activities of the compounds were performed in triplicate and are presented as the mean ± standard deviations (SD).

**Calculation of pocket volumes and preparation of structural figures**

Binding-site (pocket) volumes were caculated by using UCSF Chimera (Pettersen et al., 2004). Structural figures were prepared by using PyMol (Schrödinger LLC).

### Inhibitor synthesis

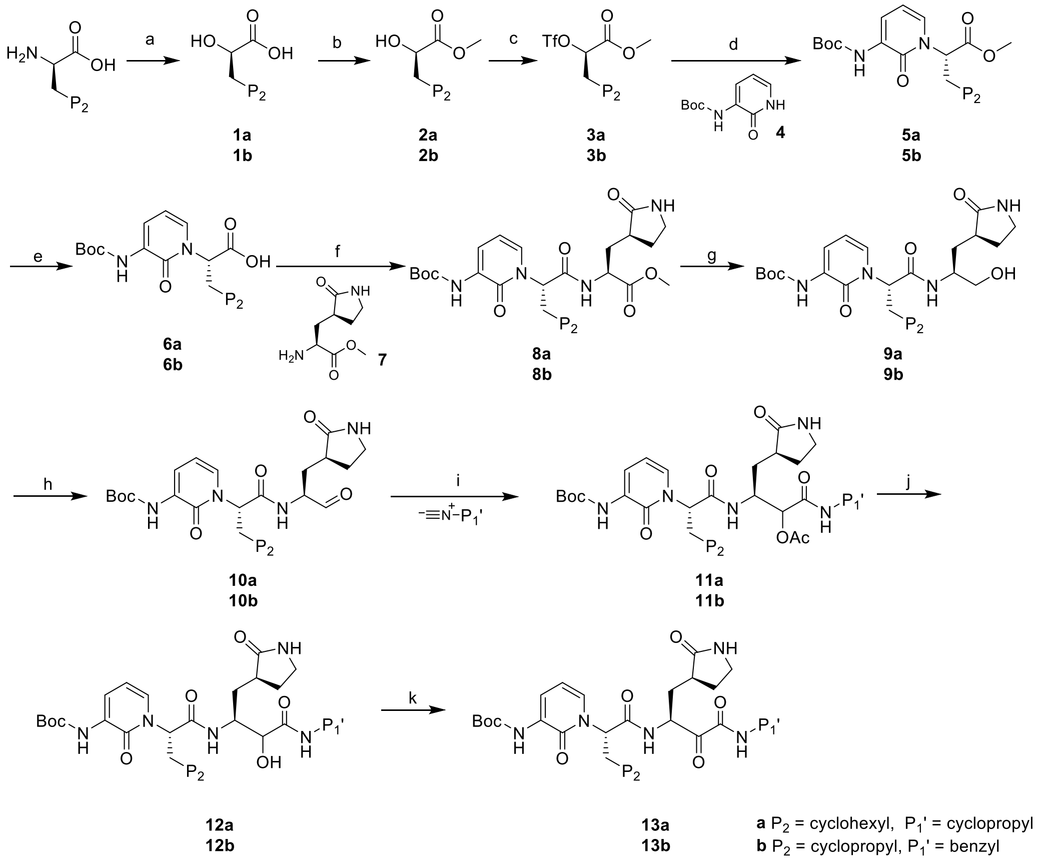


Reaction conditions: (a) $NaNO_2$, $H_2SO_4$, $H_2O$; (b) $SOCl_2$, MeOH; (c) $Tf_2O$, 2,6-lutidine, $CH_2Cl_2$; (d) NaH, THF; (e) LiOH, MeOH, $H_2O$; (f) HOBT, EDCI, $CH_2Cl_2$; (g) $NaBH_4$, MeOH; (h) DMP, $NaHCO_3$, $CH_2Cl_2$; (i) isocyanide, AcOH, $CH_2Cl_2$; (j) LiOH, MeOH, $H_2O$; (k) DMP, $NaHCO_3$, $CH_2Cl_2$

**General procedure.** Reagents were purchased from commercial sources and used without purification. HSGF 254 (0.15 - 0.2 mm thickness) was used for analytical thin-layer chromatography (TLC). All products were characterized by their NMR and MS spectra. $^1H$ NMR spectra were recorded on 300-MHz instruments. Chemical shifts are reported in parts per million (ppm, δ) down-field from tetramethylsilane. Proton coupling patterns are described as singlet (s), doublet (d), triplet (t), multiplet (m), and broad (br). Mass spectra were recorded using a Bruker ESI ion-trap HCT Ultra. HPLC spectra were recorded by an LC20A (Shimadzu Corporation) with Shim-pack GIST C18 (5 μm, 4.6x150mm) with three solvent systems (methanol/water, methanol/ 0.1% HCOOH in water, or methanol/0.1% ammonia in water). Purity was determined by reversed-phase HPLC and was ≥95% for both **13a** and **13b**.

***Synthesis of compound 1***

A solution of (*R*)-2-amino-3-cyclohexylpropanoic acid (7.74 mmol) or (*R*)-2-amino-3-cyclopropylpropanoic acid in 2N $H_2SO_4$ (15 mL) was stirred at 0 $^oC$. Then, $NaNO_2$ (5.34 g, 77.4 mmol) in $H_2O$ (6 mL) was added dropwise to the reaction. The mixture was stirred for 3 h at 0$^o$C and then allowed to warm to 20$^o$C and stirred at 20$^o$C for 16 h. The mixture was extracted with MTBE (50 mL). The combined organic phase was dried over anhydrous $Na_2SO_4$ and concentrated under vacuum to give compound **1** (as a colorless oil, 50-75% yield) without further purification.

***Synthesis of compound 2***

$SOCl_2$ (0.8 mL, 11.34 mmol) was added dropwise to the solution of compound **1** (5.72 mmol) in MeOH (20 mL) at 0$^o$C. Then the mixture was stirred for 1.5 h at 20$^o$C. The mixture was evaporated in vacuum and purified by chromatography on silica gel (PE/EA = 1/1) to give compound **2** (as colorless oil, 30-59 % yield).

M*ethyl (R)-3-cyclohexyl-2-hydroxypropanoate* **2a**:

$^{1}$H NMR (300 MHz, $CDCl_3$) δ 4.39-4.34 (m, 1H), 3.82 (s, 3H), 1.82-1.48 (m, 8H), 1.29-1.12 (m, 4H), 1.00-0.85 (m, 2H).

***Methyl (R)-3-cyclopropyl-2-hydroxypropanoate* 2b**:

$^{1}$H NMR (300 MHz, $CDCl_3$) δ 4.35 (dd, $J_1$ = 9.0 Hz, $J_2$ = 4.2 Hz, 1H), 3.80 (s, 3H), 1.81-1.72 (m, 2H), 0.92-0.68 (m, 1H), 0.50-0.43 (m, 2H), 0.15-0.05 (m, 2H).

***Synthesis of compound 3***

Compound **2** (5.32 mmol) was dissolved in DCM (10 mL) and cooled to 0$^{o}$C. 2,6-lutidine (1.5 mL, 13.26 mmol) and $Tf_2O$ (3.3 g, 11.87 mmol) were added successively. The mixture was stirred for 30 min at 0$^{o}$C. The mixture was extracted with MTBE after washing with a mixture of brine and 1N HCl (3:1 v/v), then dried over anhydrous $Na_2SO_4$, and evaporated to give compound **3** (as a brown oil, yield 82%) without further purification.

***Synthesis of compound 5***

Tert-butyl (2-oxo-1,2-dihydropyridin-3-yl)carbamate (379 mg, 1.8 mmol) was dissolved in THF (15 mL). NaH (115 mg, 2.80 mmol, 60% in oil) was added at 0$^{o}$C and then stirred for 30 min. Then compound **3** (515 mg, 1.86 mmol) in THF (10 mL) was added. The mixture was stirred for 20 h at 25 $^{o}$C. The mixture was evaporated in vacuum and purified by chromatography on silica gel (PE to EA) to give compound **5** (56-60% yield) as light yellow solid.

***Methyl (S)-2-(3-((tert-butoxycarbonyl)amino)-2-oxopyridin-1(2H)-yl)-3- cyclo-hexyl-propanoate 5a:***

$^1$H NMR (300 MHz, DMSO-$d_6$) δ 7.82-7.76 (m, 2H), 7.35 (dd, $J_1$ = 7.5 Hz, $J_2$ = 1.5 Hz, 1H), 6.30 (t, $J$ = 7.5 Hz, 1H), 5.35 (dd, $J_1$ = 11.1 Hz, $J_2$ = 4.5 Hz, 1H), 3.56 (s, 3H), 2.10-1.88 (m, 2H), 1.78-1.72 (m, 1H), 1.65-1.44 (m, 13H), 1.14-0.82 (m, 6H). ESI-MS (*m/z*): 379 $[M + H]^+$.

***Methyl (S)-2-(3-((tert-butoxycarbonyl)amino)-2-oxopyridin-1(2H)-yl)-3-cyclopropyl-propanoate 5b:***

$^1$H NMR (300 MHz, DMSO-$d_6$) δ 7.83-7.78 (m, 2H), 7.35 (dd, $J_1$ = 7.2 Hz, $J_2$ = 1.5 Hz, 1H), 6.30 (t, $J$ = 7.2 Hz, 1H), 5.36 (dd, $J_1$ = 10.8 Hz, $J_2$ = 4.5 Hz, 1H), 3.57 (s, 3H), 1.81-1.62 (m, 2H), 1.48 (s, 9H), 0.55-0.48 (m, 1H), 0.34-0.29 (m, 2H), 0.15-0.12 (m, 1H), 0.04-0.01 (m, 1H). ESI-MS (*m/z*): 337 $[M + H]^+$.

***Synthesis of compound 6***

Compound **5** (1.65 mmol) was dissolved in MeOH (15 mL) and $H_2O$ (3 mL). LiOH.H2O (139 mg, 3.31 mmol) was added. The mixture was stirred at 20°C for 1 h. Then, the mixture was adjusted to pH=6~7 with 1N HCl. Subsequently, the reaction was evaporated under vacuum and purified by chromatography on silica gel (DCM/MeOH = 10/1) to give compound **6** (452 mg, 84% yield) as light yellow solid.

***(S)-2-(3-((tert-butoxycarbonyl)amino)-2-oxopyridin-1(2H)-yl)-3-cyclohexylpropanoic acid 6a:***

$^1$H NMR (300 MHz, DMSO-$d_6$) δ 13.12 (s, 1H), 7.83-7.77 (m, 2H), 7.35 (dd, $J_1$ = 7.2 Hz, $J_2$ = 1.5 Hz, 1H), 6.30 (t, $J$ = 7.2 Hz, 1H), 5.35 (dd, $J_1$ = 10.8 Hz, $J_2$ = 4.5 Hz, 1H), 2.10-1.92 (m, 2H), 1.78-1.69 (m, 1H), 1.65-1.52 (m, 4H), 1.47 (s, 9H), 1.13-0.82 (m, 6H). ESI-MS (*m/z*): 365 [M + H]$^+$.

***(S)-2-(3-((tert-butoxycarbonyl)amino)-2-oxopyridin-1(2H)-yl)-3-cyclopropylpropanoic acid 6b:***

$^1$H NMR (300 MHz, DMSO-$d_6$) δ 13.11(s, 1H), 7.81-7.77 (m, 2H), 7.36 (dd, $J_1$ = 6.9 Hz, $J_2$= 1.5 Hz, 1H), 6.30 (t, $J$ = 6.9 Hz, 1H), 5.35 (dd, $J_1$ = 10.5 Hz, $J_2$= 4.5 Hz, 1H), 1.80-1.69 (m, 2H), 1.47 (s, 9H), 0.53-0.48 (m, 1H), 0.32-0.29 (m, 2H), 0.14-0.11 (m, 1H), 0.03-0.00 (m, 1H). ESI-MS (*m/z*): 323 [M + H]$^+$.

***Synthesis of compound 8***

HOBT (245 mg, 1.82 mmol) and EDCI (349 mg, 1.82 mmol) were added to the solution of compound **6** (1.65 mmol) in DCM (20 mL). The mixture was stirred for 1 h at 0°C. Then compound **7,** methyl (S)-2-amino-3-((S)-2-oxopyrrolidin-3-yl)propanoate (307 mg, 1.65 mmol), was added, and the mixture was adjusted to pH = 9 with $Et_3N$. The mixture was stirred at 0°C for 24 h. Then, the reaction was concentrated and purified by chromatography on silica gel (DCM/MeOH=20/1) to give compound **8** (59-67% yield) as light yellow solid.

***Methyl (S)-2-((S)-2-(3-((tert-butoxycarbonyl)amino)-2-oxopyridin-1(2H)-yl)-3-cyclohexyl propan amido)-3-((S)-2-oxopyrrolidin-3-yl)propanoate 8a:***

ESI-MS (*m/z*): 533 $[M + H]^+$

***Methyl (S)-2-((S)-2-(3-((tert-butoxycarbonyl)amino)-2-oxopyridin-1(2H)-yl)-3-cyclopropyl pro-panamido)-3-((S)-2-oxopyrrolidin-3-yl)propanoate 8b:***

$^1$H NMR (300 MHz, DMSO-$d_6$) δ 9.00-8.92 (m, 1H), 7.81-7.77 (m, 2H), 7.37-7.34 (m, 1H), 6.30 (t, $J$ = 7.2 Hz, 1H), 5.77 (dd, $J_1$ = 10.8 Hz, $J_2$ = 4.5 Hz, 1H), 4.54-4.45 (m, 1H), 3.74 (s, 3H), 3.37-3.29 (m, 2H), 2.35-2.25 (m, 2H), 1.90-1.71 (m, 5H), 1.46 (s, 9H), 0.51-0.46 (m, 1H), 0.32-0.29 (m, 2H), 0.15-0.11 (m, 1H), 0.04-0.00 (m, 1H). ESI-MS (*m/z*): 491 $[M + H]^+$.

***Synthesis of compound 9***

$NaBH_4$ (200 mg, 5.3 mmol) was added to a solution of compound **8** (0.53 mmol) in MeOH (6 mL). The mixture was stirred at 25 °C for 3 h. The reaction was concentrated and purified by chromatography on silica gel (DCM/MeOH=10/1) to give compound **9** (49%, as off-white solid).

***Tert-butyl (1-((S)-3-cyclohexyl-1-(((S)-1-hydroxy-3-((S)-2-oxopyrrolidin-3-yl)propan-2-yl) amino)-1-oxopropan-2-yl)-2-oxo-1,2-dihydropyridin-3-yl)carbamate 9a:***

ESI-MS (*m/z*): 505 $[M + H]^+$.

***Tert-butyl (1-((S)-3-cyclopropyl-1-(((S)-1-hydroxy-3-((S)-2-oxopyrrolidin-3-yl)propan-2-yl) amino)-1-oxopropan-2-yl)-2-oxo-1,2-dihydropyridin-3-yl)carbamate 9b:***

ESI-MS (*m/z*): 463 $[M + H]^+$.

***Synthesis of compound 10***

Dess-Martin Periodinane (116 mg, 0.27 mmol) and $NaHCO_3$ (8 mg, 0.09 mmol) were added to a solution of compound **9** (0.26 mmol) in DCM (15 mL). The mixture was stirred at 20°C for 1 h. The reaction was concentrated and purified by chromatography on silica gel (DCM/MeOH=20/1) to give compound **10** (83 -90% yield) as off-white solid.

***Tert-butyl (1-((S)-3-cyclohexyl-1-oxo-1-(((S)-1-oxo-3-((S)-2-oxopyrrolidin-3-yl)propan-2-yl) amino)propan-2-yl)-2-oxo-1,2-dihydropyridin-3-yl)carbamate 10a:***

ESI-MS (*m/z*): 503 $[M + H]^+$.

***Tert-butyl (1-((S)-3-cyclopropyl-1-oxo-1-(((S)-1-oxo-3-((S)-2-oxopyrrolidin-3-yl)propan-2-yl)amino)propan-2-yl)-2-oxo-1,2-dihydropyridin-3-yl)carbamate 10b:***

$^1H$ NMR (300 MHz, DMSO-$d_6$) δ 9.40 (d, $J$ = 7.8 Hz, 1H), 8.97 (dd, $J_1$ = 14.1 Hz, $J_2$= 7.2 Hz, 1H), 7.79-7.73 (m, 2H), 7.35-7.32 (m, 1H), 6.30 (t, $J$ = 7.5 Hz, 1H), 5.69-5.62 (m, 1H), 4.48-4.42 (m, 1H), 3.20-3.10 (m, 2H), 2.32-2.15 (m, 2H), 1.88-1.66 (m, 5H), 1.46 (s, 9H), 0.55-0.47 (s, 1H), 0.36-0.29 (m, 2H), 0.14-0.11 (m, 1H), 0.04-0.00 (m, 1H). ESI-MS (*m/z*): 461 $[M + H]^+$.

***Synthesis of compound 11 (general method)***

Acetic acid (26 mg, 0.44 mmol) and isocyanide (0.22 mmol) were added to the solution of compound **10** (0.22 mmol) in DCM (15 mL). The mixture was stirred at 20°C for 24 h. The reaction was concentrated and purified by chromatography on silica gel (DCM/MeOH=20/1) to give compound **11** (57-65 % yield) as off-white solid.

***Synthesis of compound 12 (general method)***

Compound **11** (0.13 mmol) was dissolved in MeOH (15 mL) and $H_2O$ (3 mL). LiOH.H2O (11 mg, 0.26 mmol) was added. The mixture was stirred at 20$^{o}$C for 20 min. The mixture was adjusted to pH=6~7 with 1N HCl. The reaction was concentrated and purified by chromatography on silica gel (DCM/MeOH=10/1) to give compound **12** (90-95 % yield) as off-white solid.

***Synthesis of compound 13 (general method)***

Compound **12** (0.115 mmol) was dissolved in DCM (15 mL). Dess-Martin periodinane (58 mg, 0.14 mmol) and $NaHCO_3$ (4 mg, 0.05mmol) were added. The mixture was stirred at 25$^{o}$C for 1 h. The reaction was concentrated and purified by chromatography on silica gel (DCM/MeOH=10/1) to give compound **13** (69-80 % yield) as off-white solid.

***Tert-butyl (1-((S)-3-cyclohexyl-1-(((S)-4-(cyclopropylamino)-3,4-dioxo-1-((S)-2-oxopyrro-lidin-3-yl)butan-2-yl)amino)-1-oxopropan-2-yl)-2-oxo-1,2-dihydropyridin-3-yl)carbamate 13a:***

$^{1}$H NMR (300 MHz, $CDCl_3$) δ 8.78-8.50 (m, 1H), 8.01-7.92 (m, 1H), 7.65 (d, *J* = 7.5 Hz, 1H), 7.10-6.90 (m, 2H), 6.35-6.14 (m, 2H), 5.85-5.75 (m, 1H), 5.25-5.10 (m, 1H), 3.24-3.13 (m, 2H), 2.75-2.71 (m, 1H), 2.49-2.20 (m, 1H), 2.10-1.81 (m, 3H) 1.80-1.52 (m, 8H), 1.49 (s, 9H), 1.25-1.01 (m, 4H), 1.00-0.73 (m, 4H), 0.56-0.51 (m, 2H). ESI-MS (*m*/*z*): 586 $[M + H]^+$.

***Tert-butyl (1-((S)-1-(((S)-4-(benzylamino)-3,4-dioxo-1-((S)-2-oxopyrrolidin-3-yl)butan-2-yl)amino)-3-cyclopropyl-1-oxopropan-2-yl)-2-oxo-1,2-dihydropyridin-3-yl)carbamate 13b:***

$^1$H NMR (300 MHz, DMSO-d$_6$) δ 9.25 (d, $J$ = 5.4 Hz, 1H), 9.00 (dd, $J_1$ = 14.1 Hz, $J_2$= 7.2 Hz, 1H), 7.79-7.69 (m, 3H), 7.35-7.22 (m, 5H), 6.30-6.24 (m, 1H), 5.69-5.61 (m, 1H), 4.97 (s, br, 1H), 4.29 (s, 2H), 3.17-3.09 (m, 2H), 2.30-2.15 (m, 2H), 1.91-1.62 (m, 5H), 1.46 (s, 9H), 0.52-0.47 (m, 1H), 0.33-0.29 (m, 2H), 0.14-0.11 (m, 1H), 0.05-0.02 (m, 1H). ESI-MS ($m/z$): 594 $[M + H]^+$.

**Determination of ADME properties**

***Plasma protein binding***. Plasma protein binding was assessed using the rapid equilibrium device (RED) system from ThermoFisher. Compounds were dissolved in DMSO. Naproxene served as control as it shows high plasma protein binding. Compounds were diluted in murine plasma (from CD-1 mice, pooled) to a final concentration of 100 µM. Dialysis buffer and plasma samples were added to the respective chambers according the manufacturer's protocol. The RED plate was sealed with a tape and incubated at 37°C for 2 hours at 800 rpm on an Eppendorf MixMate® vortex-mixer. Then samples were withdrawn from the respective chambers. To 25 µL of each dialysis sample, 25 µL of plasma and to 25 µL of plasma sample, 25 µL of dialysis buffer was added. Then 150 µL ice-cold extraction solvent (acetonitrile/$H_2O$ (90:10) containing 12.5 ng/mL caffeine as internal standard) was added. Samples were incubated for 30 min on ice. Then samples were centrifuged at 4°C at 2270 x *g* for 10 min. Supernatants were transferred to Greiner V-bottom 96-well plates and sealed with a tape. The percentage of bound compound was calculated as follows:

(1) % free = (concentration buffer chamber / concentration plasma chamber) *100

% bound = 100 % - % free.

***Plasma stability.*** **13a** dissolved in DMSO was added to mouse plasma (pH 7.4, 37°C) to yield a final concentration of 25 µM. In addition, procaine and procainamide (dissolved in DMSO) were added to mouse plasma (pH 7.4, 37°C) to yield a final concentration of 250 µM. Procaine served as positive control as it is unstable in mouse plasma. Procainamide served as negative control as it is stable in mouse plasma. The samples were incubated for 0 min, 15 min, 30 min, 60 min, 90 min, and 120 min at 37°C. At each time point, 7.5 µL of the respective sample was extracted with 22.5 µL methanol containing an internal standard for 5 min at 2000 rpm on a MixMate® vortex mixer (Eppendorf). Then samples were centrifuged for 2 min at 15.870 x *g* and the supernatants were transferred to HPLC-glass vials. 100 ng/mL glipizide was used as internal standard for HPLC-MS. Peak areas of each compound and of the internal standard were analyzed using the MultiQuant 3.0 software (AB Sciex). Peak areas of the respective compound were normalized to the internal standard peak area and to the respective peak areas at time point 0 min: (C/D)/(A/B) with A: peak area of the compound at time point 0 min, B: peak area of the internal standard at time point 0 min, C: peak area of the compound at the respective time point, D: peak area of the internal standard at the respective time point. Every experiment was repeated independently at least three times.

***Microsomal stability.*** S9 liver microsomes (mouse and human, Thermo Fisher) were thawed slowly on ice. 20 mg/mL of microsomes, 2 µL of a 100 µM solution of **13a** and 183 µL of 100 mM phosphate buffer were incubated 5 min at 37°C in a water bath. Reactions were initiated using 10 µL of 20 mM NADPH. Samples were incubated in three replicates at 37°C under gentle agitation at 150 rpm. At 0, 15, 30, and 60 min, reactions were terminated by the addition of 200 µL acetonitrile. Samples were vortexed and centrifuged at 2270 x *g* for 10 min at 4°C. The supernatants were transferred to 96-well Greiner V-bottom plates, sealed and analyzed according to the section HPLC-MS analysis below. Peak areas of the respective time point of **13a** were normalized to the peak area at time point 0 min.

***Solubility assays.*** First, a calibration curve was prepared with **13a** ranging from 2.5 to 500 µM. For determination of kinetic solubility, 0.5 mmol of **13a** was added to a 96-well plate in triplicates. 100 µL buffer of pH 7.4 was added to every replicate of **13a**. Samples were incubated at room temperature. At 15 min and 4 hrs, 10 µL were taken from every well and transferred to a new 96-well plate. Then 190 µL acetonitrile containing 12.5 ng/mL glipizide as internal standard was added. For thermodynamic solubility, 0.5 mmol of **13a** were added to a glass vial. Buffer of pH 7.4 was added and samples were incubated at 20°C for 24, 48, and 72 hrs under agitation (120 rpm). At the respective time points, 10 µL of every replicate were taken and transferred to a new 96-well plate. 190 µL acetonitrile containing 12.5 ng/mL glipizide as internal standard was added. All Samples were centrifuged for 10 min at 12°C and 2270 x *g*, supernatants were transferred to a new Greiner

96-well V-bottom plate. All samples were analyzed according to the section HPLC-MS analysis below.

***HPLC-MS analysis.*** Samples were analyzed using an Agilent 1290 Infinity II HPLC system coupled to an AB Sciex QTrap 6500plus mass spectrometer. LC conditions were as follows: column: Agilent Zorbax Eclipse Plus C18, 50x2.1 mm, 1.8 µm; temperature: 30°C; injection volume: 5 µL for plasma, 10 µL for urine, lung and BALF; 10 µL for samples from plasma stability, metabolic stability, plasma protein binding and solubility assays; flow rate: 700 µL/min; solvent A: water + 0.1 % formic acid; solvent B: 95 % acetonitrile/5 % $H_2O$ + 0.1 % formic acid; gradient: 99 % A at 0 min, 99 % A until 0.10 min, 99 % - 0% A from 0.1 min to 5.50 min, 0 % A until 6.00 min, 0 % - 99 % A from 6.00 min to 6.40 min, 99 % A until 6.50 min. Mass transitions for controls and **13a** are depicted in Table S1.

**Table S1. Mass transitions of compounds**

| | Q1 mass | Q3 mass | Time [msec] | DP [volts] | CE [volts] | CXP [volts] |
|---|---|---|---|---|---|---|
| Naproxene | 231.106 | 185.1 | 50 | 80 | 19 | 10 |
| | 231.106 | 170.2 | 50 | 80 | 33 | 12 |
| Caffeine | 195.024 | 138.0 | 30 | 80 | 25 | 14 |
| | 195.024 | 110.0 | 30 | 80 | 31 | 18 |
| **13a** | 608.228 | 508.2 | 30 | 1 | 39 | 28 |
| | 608.228 | 313.1 | 30 | 1 | 53 | 34 |
| Glipizide | 444.027 | 318.9 | 30 | -200 | -38 | -12 |
| | 444.027 | 170.0 | 30 | -200 | -38 | -12 |
| Procaine | 235.744 | 163.0 | 30 | 80 | 21 | 18 |
| | 235.744 | 120.0 | 30 | 80 | 39 | 12 |
| Procainamide | 236.773 | 100.0 | 30 | 80 | 21 | 12 |
| | 236.773 | 120.0 | 30 | 80 | 31 | 14 |

**Determination of pharmacokinetic properties**

***Mice.*** For pharmacokinetic experiments, outbred male CD-1 mice (Charles River, Germany), 4 weeks old, were used. The animal studies were conducted in accordance with the recommendations of the European Community (Directive 86/609/EEC, 24 November 1986). All animal procedures were performed in strict accordance with the German regulations of the Society for Laboratory Animal Science (GV-SOLAS) and the European Health Law of the Federation of Laboratory Animal Science Associations (FELASA). Animals were excluded from further analysis if sacrifice was necessary according to the human endpoints established by the ethical board. All experiments were approved by the ethical board of the Niedersächsisches Landesamt für Verbraucherschutz und Lebensmittelsicherheit, Oldenburg, Germany (LAVES; permit no. 33.19-42502-04-15/1857).

***Pharmacokinetic (PK) study.*** **13a** was dissolved in olive oil and water containing lecithin (40 mg lecithin dissolved in 800 µL of water). Mice were administered **13a** subcutaneously (s.c.) at 20 mg/kg. About 20 µl of whole blood was collected serially from the lateral tail vein at time points 0.25, 0.5, 1, 2, 4, and 8 h post administration. After 24 h, mice were sacrificed and blood was collected from the heart. Whole blood was collected into Eppendorf tubes coated with 0.5 M EDTA and immediately spun down at 15870 x *g* for 10 min at 4°C. The plasma was transferred into a new Eppendorf tube and then stored at −80°C until analysis.

***PK sample preparation and analysis.*** First, a calibration curve was prepared by spiking different concentrations of **13a** into mouse plasma, mouse urine, murine broncheo-alveolar lavage fluid (BALF), or homogenized lung from CD-1 mice. Caffeine was used as an internal standard. In addition, quality control samples (QCs) were prepared for **13a** in plasma, urine, BALF, and homogenized lung. The following extraction procedures were used: 7.5 µL of a plasma sample or 10 µL of a urine sample (calibration samples, QCs or PK samples) was extracted with 25 µL of acetonitrile containing 12.5 ng/mL of caffeine as internal standard for 5 min at 2000 rpm on an Eppendorf MixMate® vortex mixer; 50 µL of a lung sample (concentration adjusted to 50 mg/mL, calibration samples, QCs or PK samples) was extracted with 50 µL of acetonitrile containing 12.5 ng/mL caffeine as internal standard for 5 min at 2000 rpm on an Eppendorf MixMate® vortex mixer. BALF samples were prepared as follows: 300 µL methanol were added to 200 µL of a BALF sample (calibration samples, QCs or PK samples) and samples were evaporated to dryness in an Eppendorf concentrator. Then, 80 µL of water were added to every dry sample and incubated for 10 min at 2000 rpm on an Eppendorf MixMate® vortex mixer. Then, 20 µL of acetonitrile containing 125 ng/mL caffeine were added and samples incubated for another 5 min at 2000 rpm on an Eppendorf MixMate® vortex mixer. Then samples were spun down at 15870 x *g* for 10 min. Supernatants were transferred to standard HPLC-glass vials as described in the section HPLC-MS analysis above. Peaks of PK samples were quantified using the calibration curve. The accuracy of the calibration curve was determined using QCs independently prepared on different days. PK parameters were determined using a non-compartmental analysis with PKSolver (Zhang et al., 2010).

**Supplementary Tables**

**Table S2. Diffraction data and model refinement statistics**

| **Protein / Ligand** | **$M^{pro}$ free enzyme** | **$M^{pro}$ + 13b (monoclinic form)** | **$M^{pro}$ + 13b (orthorhombic form)** |
|---|---|---|---|
| **PDB entry** | 6Y2E | 6Y2F | 6Y2G |
| **Data collection statistics** | | | |
| X-ray source | BESSY 14.2 | BESSY 14.2 | BESSY 14.2 |
| Wavelength [Å] | 0.9184 | 0.9184 | 0.9184 |
| $V_m$ [$A^3$/Da] | 2.01 | 2.75 | 2.67 |
| Solvent content [%] | 38.8 | 55.3 | 54.0 |
| Space group | *C*2 | *C*2 | $P2_12_12_1$ |
| Unit cell dimensions [Å] | *a* = 114.98, *b* = 53.76, *c* = 44.77 | *a* = 98.08, *b* = 80.93, *c* = 51.66 | *a* = 68.57, *b* = 101.60, *c* = 103.70 |
| Unit cell dimensions [º] | α = γ = 90, β = 101.24 | α = γ = 90, β = 114.84 | α = β = γ = 90 |
| Resolution range [a] [Å] | 48.53 - 1.75 (1.84 - 1.75) | 43.37 - 1.95 (2.06 - 1.95) | 41.39 - 2.20 (2.32 - 2.20) |
| Number of observations | 185,991 (27,913) | 182,301 (27,374) | 492,459 (64,837) |
| Number of unique reflections | 27,173 (3,926) | 26,757 (3,919) | 37,519 (5356) |
| Completeness [%] | 100.0 (100.0) | 100.0 (100.0) | 100.0 (100.0) |
| Mean I/σ(I) | 15.0 (2.1) | 15.5 (2.2) | 13.1 (2.1) |
| Multiplicity | 6.8 (7.1) | 6.8 (7.0) | 13.1 (12.1) |
| $R_{merge}$ [b] [%] | 0.082 (0.967) | 0.076 (0.826) | 0.170 (1.388) |
| $R_{pim}$ [c] [%] | 0.034 (0.387) | 0.032 (0.335) | 0.048 (0.414) |
| $CC_{1/2}$ [d] | 0.999 (0.747) | 0.999 (0.809) | 0.998 (0.662) |
| Wilson *B*-factor [$A^2$] | 23.0 | 31.2 | 35.5 |
| **Refinement statistics** | | | |
| $R_{cryst}$ [e] /$R_{free}$ [f] [%] | 17.12/22.24 | 17.79/21.91 | 18.62/24.73 |
| r.m.s.d. in bond lengths [Å] | 0.009 | 0.010 | 0.010 |
| r.m.s.d. in bond angles [°] | 1.566 | 1.728 | 1.815 |
| Clashscore [g] | 2 | 3 | 4 |

| | | | |
|---|---|---|---|
| Average B-factor for protein atoms [Å$^2$] | 26.71 | 39.00 | 44.17 |
| Average B-factor for ligand atoms [Å$^2$] | N/A | 47.24 | 49.36 |
| Average B-factor for water molecules [Å$^2$] | 36.74 | 47.30 | 43.79 |
| Number of protein atoms | 2373 | 2342 | 4675 |
| Number of ligand atoms | N/A | 43 | 86 |
| Number of water molecules | 313 | 177 | 313 |
| **Ramachandran plot** | | | |
| Preferred regions [%] | 96.37 | 94.32 | 93.83 |
| Allowed regions [%] | 3.30 | 5.35 | 6.17 |
| Outlier regions [%] | 0.33 | 0.33 | 0 |

[a] The highest resolution shell is shown in parantheses.

[b] $R_{merg} = \sum_{hkl}\sum_{i=1}^{n}|I_i(hkl) - \bar{I}(hkl)|/\sum_{hkl}\sum_{i=1}^{n} I_i(hkl)$

[c] $R_{pim} = \sum_{hkl}\sqrt{1/(n-1)}\sum_{i=1}^{n}|I_i(hkl) - \bar{I}(hkl)|/\sum_{hkl}\sum_{i=1}^{n} I_i(hkl)$ (Weiss & Hilgenfeld, 1997)

[d] $CC_{1/2}$ is the correlation coefficient determined by two random half data sets (Karplus & Diederichs, 2012)

[e] $R_{cryst} = \sum_{hkl}|F_o(hkl) - F_c(hkl)|/\sum_{hkl}|F_o(hkl)|$

[f] $R_{free}$ was calculated for a test set of reflections (5%) omitted from the refinement.

[g] Clashscore is defined as the number of clashes calculated for the model per 1000 atoms (including hydrogens) of the model. Hydrogens were added by MolProbity (Chen et al., 2010).

**Table S3. Pharmacokinetic parameters of 13a after subcutaneous administration (20 mg/kg)**

| | **20 mg/kg SC** |
|---|---|
| **t1/2 [h]** | 0.98 ± 0.1 |
| **Tmax [h]** | 0.42 ± 0.1 |
| **Cmax [ng/mL]** | 334.50 ± 109.2 |
| **AUC 0-t [ng/mL*h]** | 551.20 ± 67.7 |
| **MRT [h]** | 1.59 ± 0.2 |
| **Vz/F [l/kg]** | 48.57 ± 7.7 |
| **Cl/F [mL/kg/min]** | 565.59 ± 61.0 |

t1/2: half-life, Tmax: time point at which maximal concentration is reached, Cmax: maximal concentration, AUC 0-t: area under the cure from time point 0 until t, MRT: mean residence time, Vz: volume of distribution, Cl: clearance, F: fraction/bioavailability

**References and Notes:**


Chen V. B., Arendall W. B., Headd J. J., Keedy D. A., Immormino R. M., Kapral G. J., Murray L. W., Richardson J. S., Richardson D. C., *MolProbity*: all-atom structure validation for macromolecular crystallography. *Acta Crystallogr. D Biol. Crystallogr*. **66**, 12-21 (2010).

Emsley P., Lohkamp B., Scott W. G., Cowtan K., Features and development of Coot. *Acta Crystallogr. D Biol. Crystallogr.* **66**, 486-501 (2010).

Evans P. R., Scaling and assessment of data quality. *Acta Crystallogr. D Biol. Crystallogr.* **62**, 72-82 (2006).

Evans P. R., An introduction to data reduction: space-group determination, scaling and intensity statistics. *Acta Crystallogr. D Biol. Crystallogr.* **67**, 282-292 (2011).

Karplus P. A., Diederichs K., Linking crystallographic model and data quality. *Science* **336**, 1030-1033 (2012).

Krug M., Weiss M. S., Heinemann U., Mueller U., XDSAPP: a graphical user interface for the convenient processing of diffraction data using XDS. *J. Appl. Crystal.* **45**, 568-572 (2012).

Lebedev A. A., Young P., Isupov M. N., Moroz O. V., Vagin A. A., Murshudov G. N., JLigand: a graphical tool for the CCP4 template-restraint library. *Acta Crystallogr. D Biol. Crystallogr.* **68**, 431-440 (2012).

Mueller U., Darowski N., Fuchs M. R., Förster R., Hellmig M., Paithankar K. S., Pühringer S., Steffien M., Zocher G., Weiss M. S., Facilities for macromolecular crystallography at the Helmholtz-Zentrum Berlin. *J. Synchrotron Radiat.* **19**, 442-449 (2012).

Murshudov G. N., Skubak P., Lebedev A. A., Pannu N. S., Steiner R. A., Nicholls R. A., Winn M. D., Long F., Vagin A. A., REFMAC5 for the refinement of macromolecular crystal structures. *Acta Crystallogr. D Biol. Crystallogr.* **67**, 355-367 (2011).

Pettersen E. F., Goddard T. D., Huang C. C., Couch G. S., Greenblatt D. M., Meng E. C., Ferrin T. E., UCSF Chimera - a visualization system for exploratory research and analysis. *J. Comput. Chem.* **25**, 1605-1612 (2004).

Vagin A., Teplyakov A., Molecular replacement with MOLREP. *Acta Crystallogr. D Biol. Crystallogr.* **66**, 22-25 (2010).

Weiss M. S., Hilgenfeld R., On the use of the merging *R* factor as a quality indicator for X-ray data. *J. Appl. Cryst.* **30**, 203-205 (1997).

Winn M., Ballard C. C., Cowtan K. D., Dodson E. J., Emsley P., Evans P. R., Keegan R. M., Krissinel E. B., Leslie A. G., McCoy A., McNicholas S. J., Murshudov G. N., Pannu N. S.,

Potterton E. A., Powell H. R., Read R. J., Vagin A., Wilson K. S., Overview of the CCP4 suite and current developments. *Acta Crystallogr. D Biol. Crystallogr*. **67**, 235-242 (2011).

Xue X., Yang H., Shen W., Zhao Q., Li J., Yang K., Chen C., Jin Y., Bartlam M., Rao Z., Production of authentic SARS-CoV $M^{pro}$ with enhanced activity: application as a novel tag-cleavage endopeptidase for protein overproduction. *J. Mol. Biol.* **366,** 965-975 (2007).

Zhang Y., Huo M., Zhou J., Xie S., PKSolver: An add-in program for pharmacokinetic and pharmacodynamic data analysis in Microsoft Excel. *Comput. Methods Programs Biomed.* **99,** 306–314 (2010).

Zhu L., George S., Schmidt M. F., Al-Gharabli S. I., Rademann J., R. Hilgenfeld, Peptide aldehyde inhibitors challenge the substrate specificity of the SARS-coronavirus main protease. *Antiviral Res.* **92**, 204-212 (2011).
